# Host subversion of bacterial metallophore usage drives copper intoxication

**DOI:** 10.1101/2023.05.30.542972

**Authors:** Saika Hossain, Jacqueline R. Morey, Stephanie L. Neville, Katherine Ganio, Jana N. Radin, Javiera Norambuena, Jeffrey M. Boyd, Christopher A. McDevitt, Thomas E. Kehl-Fie

## Abstract

Microorganisms can acquire metal ions in metal-limited environments using small molecules called metallophores. While metals and their importers are essential, metals can also be toxic, and metallophores have limited ability to discriminate metals. The impact of the metallophore-mediated non-cognate metal uptake on bacterial metal homeostasis and pathogenesis remains to be defined. The globally significant pathogen *Staphylococcus aureus* uses the Cnt system to secrete the metallophore staphylopine in zinc-limited host niches. Here, we show that staphylopine and the Cnt system facilitate bacterial copper uptake, potentiating the need for copper detoxification. During *in vivo* infection, staphylopine usage increased *S. aureus* susceptibility to host-mediated copper stress, indicating that the innate immune response can harness the antimicrobial potential of altered elemental abundances in host niches. Collectively, these observations show that while the broad-spectrum metal-chelating properties of metallophores can be advantageous, the host can exploit these properties to drive metal intoxication and mediate antibacterial control.

**IMPORTANCE:** During infection bacteria must overcome the dual threats of metal starvation and intoxication. This work reveals that the zinc-withholding response of the host sensitizes *Staphylococcus aureus* to copper intoxication. In response to zinc starvation *S. aureus* utilizes the metallophore staphylopine. The current work revealed that the host can leverage the promiscuity of staphylopine to intoxicate *S. aureus* during infection. Significantly, staphylopine-like metallophores are produced by a wide range of pathogens, suggesting that this is a conserved weakness that the host can leverage to toxify invaders with copper. Moreover, it challenges the assumption that the broad-spectrum metal binding of metallophores is inherently beneficial to bacteria.

## INTRODUCTION

Transition metals are essential for all forms of life. However, their bioavailability is limited in both the environment and host niches infected by pathogens. To overcome metal starvation, microbes from all three domains of life produce small metal-binding molecules or metallophores^1–3^. While initially thought to selectively import a single metal, recent advances have revealed broader metal-binding and import capabilities^4,5^. Given the frequent restricted bioavailability of multiple essential metals in the environment and host, the broad-spectrum metal-binding of metallophores is generally regarded as beneficial^5,6^. However, despite the essentiality of transition metals, they can also mediate toxicity^7^. Accordingly, microbes tightly regulate the expression of metal uptake systems by inducing their expression as the abundance of their target metal decreases to prevent starvation and avoid intoxication in metal-replete environments^8^. Metallophore synthesis is also induced by the absence of select metals^9,10^. This leads to the question of whether the promiscuity of metallophore metal recruitment is beneficial or detrimental. Metallophore-derived therapeutics and metal-derived strategies for the environmental control of microbial populations are being increasingly used^11^. Understanding how these small molecules influence metal homeostasis at the host-pathogen interface and microbial survival is necessary to advance human health and environmental engineering.

Immunological proteins such as transferrin, lactoferrin, and calprotectin reduce the availability of essential elements, including manganese (Mn), iron (Fe), and zinc (Zn) at sites of infection in an attempt to starve invaders^12,13^. To overcome metal limitation, pathogens express a variety of metal uptake systems, including metallophores and their cognate importers^14,15^. The disruption of metallophore synthesis or import can impair the ability of diverse Gram-positive and Gram-negative pathogens to compete with the host for Fe and/or Zn, thereby reducing virulence^10,16^. In addition to starvation, pathogens also encounter host-mediated metal intoxication during infection, with the host actively employing elements such as copper (Cu) and Zn to prevent infection^7,17^. Although the precise routes for metal ion influx within host niches remain to be fully defined, recent studies have shown that this can occur at both the tissue and cellular level. For example, Cu accumulation within the phagolysosome of phagocytic cells has been shown to potentiate the killing of invading microbial pathogens^18–20^. Resistance to metal intoxication generally requires a combination of regulatory and stress response systems that frequently involve the expression of dedicated metal efflux pumps. Accordingly, loss of dedicated Cu efflux pathways frequently attenuates the ability of pathogenic bacteria to resist phagocytic killing or cause infection^19–22^. However, while pathogenic bacteria and other microbes encounter Cu intoxication, the molecular pathways that enable unregulated Cu access to the cytosol remain poorly defined.

Here, we investigated the hypothesis that broad-spectrum metal-binding metallophores render microbes susceptible to Cu intoxication using a recently described family of opine metallophores that are encoded by pathogenic and environmental organisms from multiple genera, including *Staphylococcus, Pseudomonas, Yersinia, Paenibacillus, Serratia, Bacillus,* and *Vibrio*^2,3^. Although the characterized members of the family are regulated by Zn availability, these metallophores have limited metal specificity^9,10,15^. Staphylopine (StP), the archetypal opine metallophore, is produced by the globally significant pathogen *Staphylococcus aureus* in response to Zn limitation^15^. *S. aureus* colonizes about 30% of the world’s population and is a major cause of antibiotic-resistant infections^23^. StP and its cognate importer are the primary mechanism used by *S. aureus* to compete with the host for Zn^10^. StP is produced by CntKLM, exported by CntE, and reimported in the metal-complex form by CntABCDF^15,24^. *S. aureus* also employs a Zn-specific ABC family transporter, AdcABC, that recruits Zn cations via cell-associated protein components^10,25^. Nevertheless, StP and the Cnt system are necessary for infection^10^. Here, we examined the role of Zn limitation and the StP-Cnt system on *S. aureus* susceptibility to Cu intoxication.

## RESULTS

### Zinc limitation increases activation of the copper stress response

If metallophores contribute to Cu accumulation within the bacterial cytosol, it follows that the Cu stress response will be activated in environments that trigger the production of small opine molecules. In *S. aureus,* the primary mechanism of Cu tolerance is the Cu(I)-specific efflux pump CopA, which is induced in response to cytoplasmic accumulation of this metal^26^. Following growth in the presence of Zn, significant *copA* induction, assessed using a fluorescent reporter, required concentrations of CuSO_4_ ≥250 μM (Fig. 1A). In contrast, in medium rendered Zn-deplete via chelex treatment^27^, significant *copA* induction occurred at concentrations as low as 230 nM CuSO_4_ (Fig. 1B). This indicates that three orders of magnitude less Cu is necessary to induce the Cu stress response in Zn-limited conditions than in Zn-replete conditions. Upon exposure to Zn limitation, both the Adc and StP-Cnt systems are induced in *S. aureus*^10^. To determine whether activation of the Cu stress response was driven by either system, *copA* expression was determined in Δ*adcA,* Δ*cntA*, and Δ*cntKLM*. AdcA and CntA are the solute-binding proteins for their respective transporters and are necessary for function. Thus, the absence of CntA renders *S. aureus* dependent on the Adc system, while the loss of AdcA necessitates the use of the StP-Cnt system^10,24^. Loss of CntA, or CntKLM, but not AdcA, significantly ablated the expression of *copA* in Zn-deplete medium (Fig. 1C). Similar to wild type, supplementation with 10 μM ZnSO_4_ abrogated the induction of *copA* in Δ*adcA* (Fig. 1D). The latter observation was unexpected, as Δ*adcA* is reliant upon the StP-Cnt system to obtain Zn. This observation is explained by subsequent analyses that revealed that the *cnt* operon remained Zn-responsive in Δ*adcA* (Supplemental Fig. 1). These observations indicate that activation of the Cu stress response is predominately dependent on the production and import of StP by the Cnt system.

**Figure 1.**
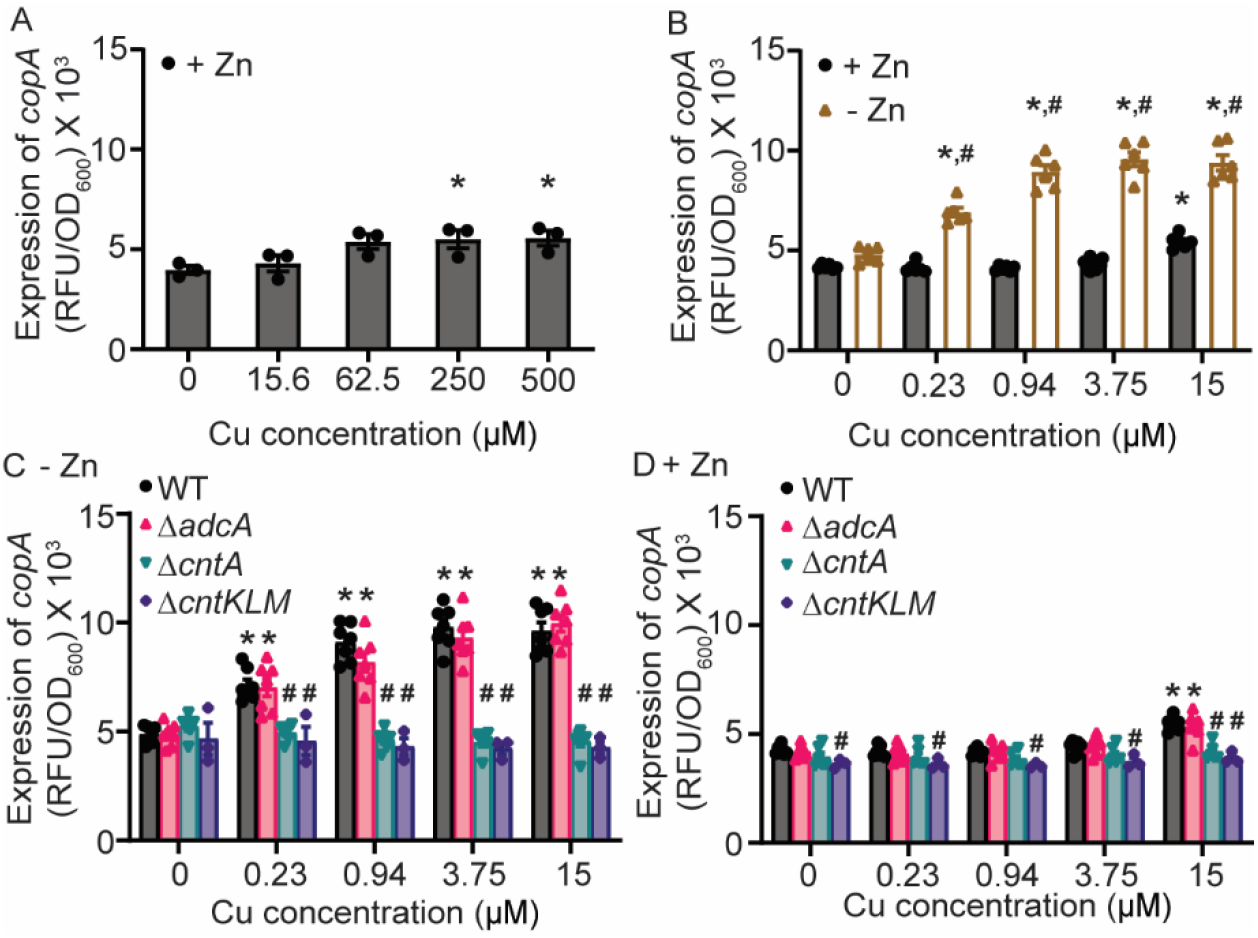
Zinc limitation increases activation of the copper stress response. (A-D) *S. aureus* Newman wild type and the indicated strains containing P*_copA_*-YFP reporter were grown in NRPMI containing a range of CuSO_4_ concentrations in the presence or absence of 10 μM ZnSO_4_ as specified. The strains were precultured (A) with or (B-D) without 10 μM ZnSO_4_. The expression of *copA* was assessed by measuring fluorescence at T = 6. * = p ≤ 0.05 relative to the same condition or strain without Cu via (A) One-way ANOVA with Dunnett’s posttest, (B) two-way ANOVA with Sidak’s posttest, or (C & D) two-way ANOVA with Dunnett’s posttest. (B) # = p ≤ 0.05 relative to bacteria grown in the presence of Zn at the same concentration of Cu via two-way ANOVA with Sidak’s posttest. (C & D) # = p ≤ 0.05 relative to wild type bacteria at the same Cu concentration via two-way ANOVA with Dunnett’s posttest. n ≥ 3. Error bars = SEM.

### Metallophore usage increases sensitivity to copper toxicity

Next, the potential role of the Cnt system in driving Cu poisoning in *S. aureus* was investigated. Wild type *S. aureus* Newman and derivatives lacking AdcA and CntA were grown in the presence and absence of Cu in a Zn-limited medium. In the absence of Cu, all strains showed similar growth (Fig. 2A). Upon supplementation with CuSO_4_ up to 1000 μM, Δ*cntA* did not show significantly impaired growth (Fig. 2B-E), indicating that AdcA-dependent Zn acquisition does not increase sensitivity to extracellular Cu. In contrast, Δ*adcA* showed substantially and statistically significant reduced growth relative to wild type and Δ*cntA* (Fig. 2D & E). Plasmid-based expression of AdcA (Fig. 2F) or supplementation with excess ZnSO_4_ (Supplemental Fig. 2) ablated the growth impairment. To mimic the sequential Zn limitation followed by Cu exposure experienced during infection, the strains were grown in a Zn-limited medium and then spot-plated onto a solid medium with and without CuSO_4_. In the absence of Cu, wild type, Δ*adcA,* and Δ*cntA* were recovered equally. However, in the presence of 25 or 50 μM CuSO_4_ supplementation, Δ*adcA* showed a 10-fold greater sensitivity to Cu than wild type or Δ*cntA* in 60% (3/5) and 100% (5/5) of assays respectively (Fig. 2G, Supplemental Fig. 3A). To this point, a methicillin-sensitive strain that possesses a single Cu efflux pump was used^28^. To evaluate if the use of the Cnt-StP system renders strains with multiple Cu efflux systems sensitive to Cu intoxication, the methicillin-resistant isolate USA300 LAC was assessed. Similar to Newman, USA300 LAC lacking AdcA was more sensitive to Cu exposure than wild type (Supplemental Fig. 3B). These findings indicate that during conditions of Zn limitation, the use of StP renders *S. aureus* more vulnerable to Cu intoxication even in the presence of multiple functional detoxification system.

**Figure 2.**
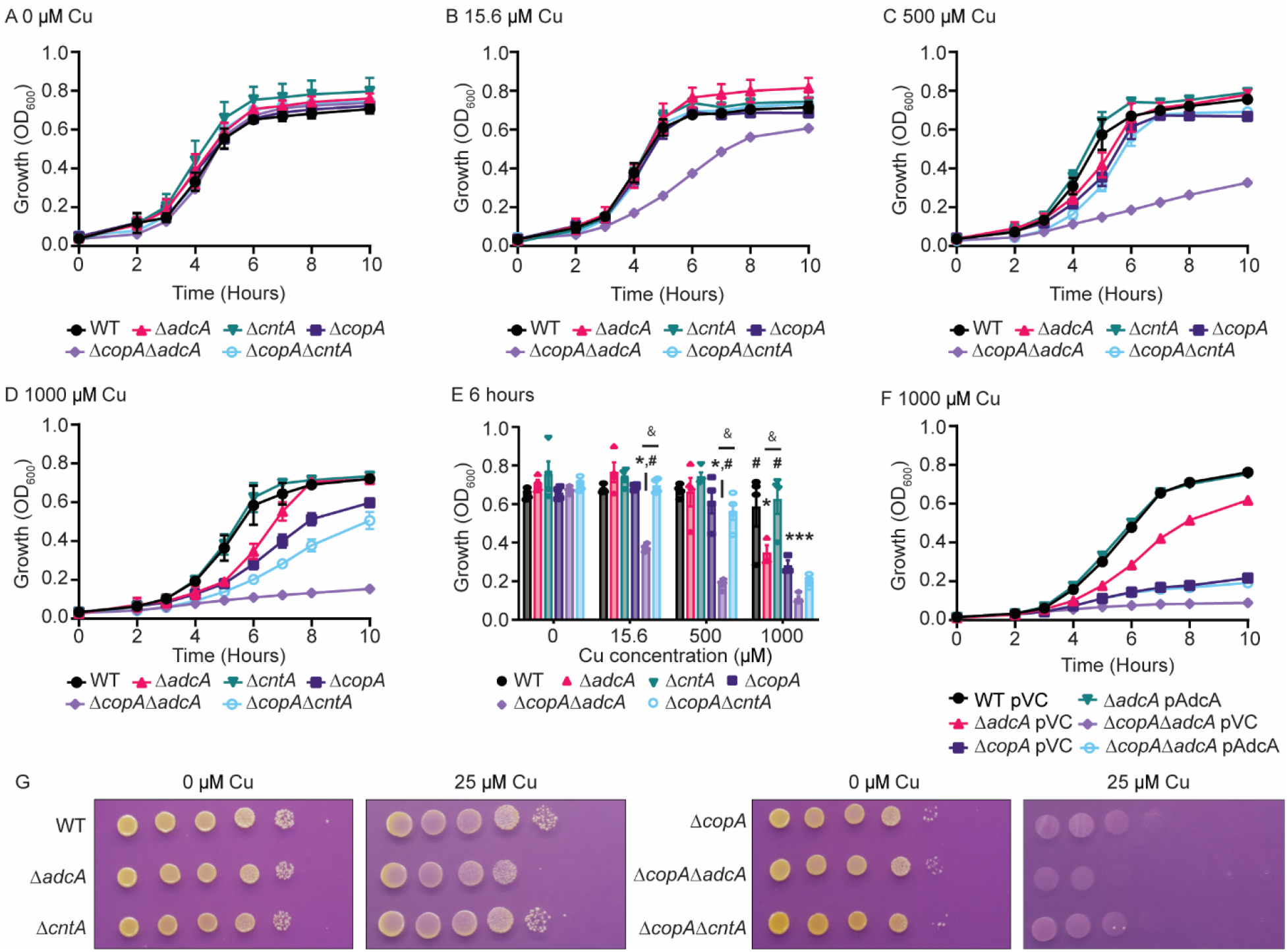
Metallophore import increases the sensitivity of *S. aureus* to copper toxicity. (A-E) *S. aureus* Newman wild type and the indicated strains were grown in Zn-limited NRPMI supplemented with CuSO_4_ as specified. (F) *S. aureus* Newman wild type and the indicated mutants carrying either vector control (pVC) or a plasmid expressing AdcA (pAdcA) were grown in Zn-limited NRPMI medium containing 1000 µM CuSO_4_. (A-F) Growth was assessed by measuring absorbance at OD_600_ over time. Statistical analysis is presented in panel E showing growth of the indicated strains at T = 6 hours. * = p ≤ 0.05 relative to wild type bacteria at the same Cu concentration via two-way ANOVA with Tukey’s posttest. # = p ≤ 0.05 relative to Δ*copA* at the same concentration via two-way ANOVA with Tukey’s posttest. & = p ≤ 0.05 for the indicated comparison via two-way ANOVA with Tukey’s posttest. n ≥ 3. Error bars indicate SEM. (G) *S. aureus* Newman wild type, and the indicated mutants were cultured in Zn-limited NRPMI, spot plated onto plates with or without Cu as mentioned. n = 5. Representative images of the spot plates are shown.

### Broad-spectrum metal-binding sensitizes bacteria to multiple antimicrobial activities associated with metal intoxication

Copper toxicity in bacteria can arise by direct modalities of action or indirectly through the disruption of essential metal uptake^29,30^. To evaluate which modes of action occurred in *S. aureus*, metal accumulation was assessed in wild type, Δ*adcA*, and Δ*cntA* following growth in Zn-deplete medium supplemented with 0, 15, or 500 μM CuSO_4_ (Fig. 3A-F). In the absence of Cu supplementation, no difference in metal content was observed between the three strains, with the exception of ^63^Cu (Fig. 3A, B, Supplemental Fig. 4). Loss of CntA was associated with a significant reduction in ^63^Cu showing the contribution of StP to *S. aureus* Cu accumulation. With 15 μM CuSO_4, 63_Cu accumulation increased in wild type and Δ*adcA* by ∼50-fold (Fig. 3C). In this condition, Δ*cntA* accumulated ∼8-fold less ^63^Cu than wild type or Δ*adcA*, indicating that the StP-Cnt system is the primary driver of Cu accumulation. In the presence of 500 μM CuSO_4_, wild type, Δ*adcA,* and Δ*cntA* accumulated similar levels of cellular ^63^Cu (Fig. 3E). This suggests that other import mechanisms exist but that high levels of Cu are necessary to drive Cu uptake via such means. Strains lacking AdcA also accumulated less Zn in this condition (Fig. 3F). No other change in metal content among these three strains was observed with 15 μM or 500 μM CuSO_4_ (Supplemental Fig. 4).

**Figure 3.**
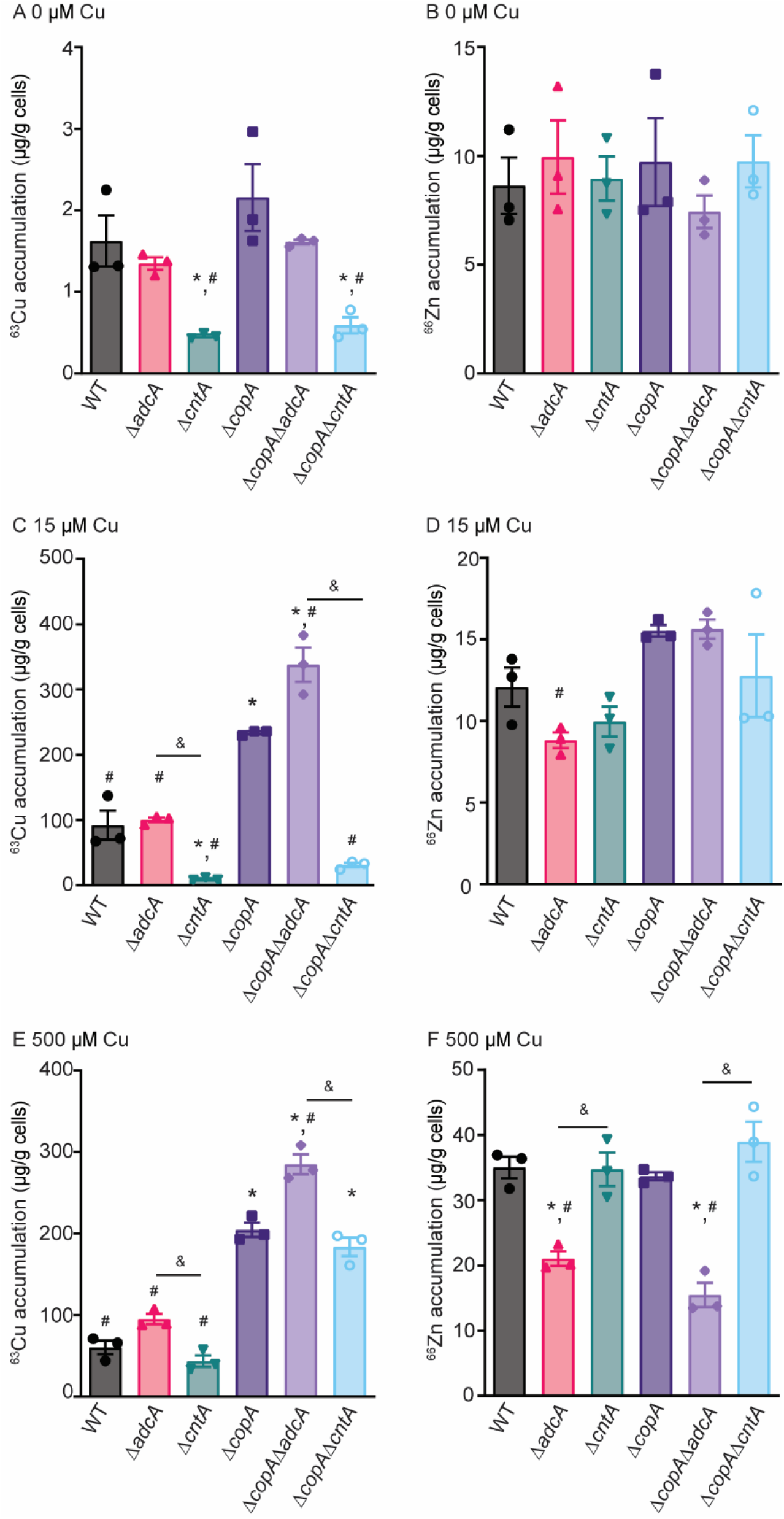
The Cnt system leads to increased Cu accumulation in *S. aureus* in Zn-limited conditions. *S. aureus* Newman wild type and the indicated mutants were grown in Zn-limited medium supplemented with (A, B) 0 μM, (C, D) 15 μM, and (E, F) 500 μM CuSO_4_, and (A, C, E) ^63^Cu or (B, D, E) ^66^Zn content was assessed using ICP-MS. * = p < 0.05 via one-way ANOVA relative to wild type bacteria using Tukey’s posttest. # = p <0.05 via one-way ANOVA relative to Δ*copA* using Tukey’s posttest. & = p < 0.05 via one-way ANOVA for the indicated comparison via Tukey’s posttest. n = 3. Error bars indicate SEM.

The accumulation data suggests that in Zn-limited environments, Cu could both disrupt cellular processes and prevent Zn uptake. Therefore, studies were taken to better understand both potential impacts. Within the cytosol, Cu can associate with proteins and potentially disrupt their function. In *S. aureus*, Cu accumulation has been shown to reduce the activity of glyceraldehyde-3-phosphate dehydrogenase (GAPDH)^29^. To evaluate if the observed intracellular Cu disrupts cellular processes, GAPDH activity was assessed in wild type, Δ*adcA,* and Δ*cntA*. In the absence of Cu, GAPDH activity is comparable across all strains tested (Fig. 4A). In the presence of 15 μM CuSO_4_ Δ*adcA* had reduced activity relative to Δ*cntA* in the same condition or to wild type and itself in unsupplemented medium indicating that the observed Cu accumulation negatively impacts cellular processes.

**Figure 4.**
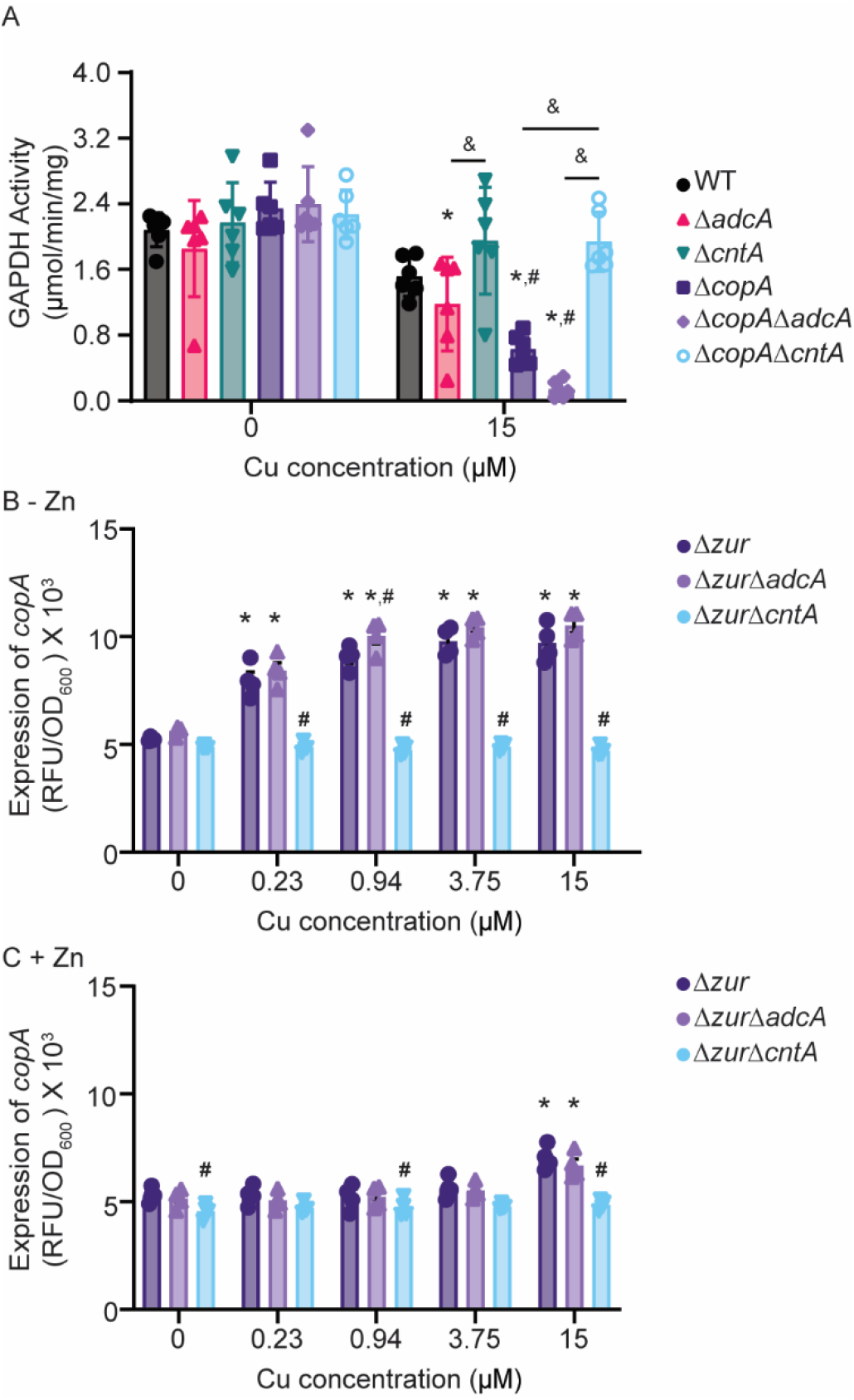
The Cnt system sensitizes *S. aureus* to Cu intoxication. (A) *S. aureus* Newman wild type and the indicated strains were grown in Zn-limited NRPMI medium supplemented with CuSO_4_ as specified and glyceraldehyde-3-phosphate dehydrogenase (GAPDH) activity was assessed. * = p ≤ 0.05 relative to wild type bacteria and the same strain grown in the absence of Cu via two-way ANOVA with Sidak’s posttest. # = p ≤ 0.05 relative to wild type bacteria grown in the same concentration of Cu via two-way ANOVA for the indicated comparison via Tukey’s posttest. & = p < 0.05 via two-way ANOVA for the indicated comparison via Tukey’s posttest. N = 6. Error bars = SEM. *S. aureus* Newman Δ*zur,* Δ*zur*Δ*adcA* and Δ*zur*Δ*cntA* carrying the P*_copA_*-YFP reporter were grown in (B) Zn-deplete and (C) Zn-replete medium in the presence and absence of Cu. The expression of *copA* was assessed by measuring fluorescence at T = 6. * = p ≤ 0.05 relative to the same strain in the absence of Cu via two-way ANOVA with Dunnett’s posttest. # = p ≤ 0.05 relative to Δ*zur* at the same Cu concentration via two-way ANOVA with Dunnett’s posttest n = 3. Error bars = SEM.

Supplementation with 500 µM CuSO_4_, but not 15 µM, resulted in Δ*adcA* accumulating less ^66^Zn than wild type or Δ*cntA* (Fig. 3F). These data suggest that high levels of Cu may interfere with StP-mediated Zn uptake. To further explore this inference, *copA* expression was assessed in strains lacking the zinc uptake regulator, Zur, which results in constitutive expression of both the Adc and Cnt-StP systems^10^. In the absence of Zn supplementation, expression of *copA* was induced by ≥230 nM CuSO_4_ supplementation in the *S. aureus* Δ*zur* and Δ*zur*Δ*adcA* mutants (Fig. 4B). Upon supplementation with 10 μM ZnSO_4_, *copA* induction was muted and could only be observed when CuSO_4_ supplementation exceeded 15 μM (Fig. 4C). In the Δ*zur*Δ*cntA* mutant, no increase in *copA* induction was observed in either the presence or absence of Zn at any CuSO_4_ concentration tested (Fig. 4B-C). These observations show that Cu and Zn availability influences the uptake of each other, albeit in a manner that is predominantly dependent upon their route of import, i.e. Adc vs. StP-Cnt. This inference is consistent with StP having the ability to bind and facilitate the import of both metal ions. Cumulatively, these results show that Zn limitation sensitizes *S. aureus* to two modes of Cu intoxication, with low Cu concentrations sufficient to prompt toxic import by StP, while high levels drive Cu import and hinder the acquisition of Zn.

### Copper efflux systems are necessary to mitigate the impact of metallophore usage

The induction of *copA* in response to nanomolar levels of extracellular Cu in Zn-limited environments suggests that the risk of metallophore-mediated metal uptake is potentially mitigated by the Cu stress response. This inference was assessed by measuring the sensitivity of *S. aureus* Δ*copA*, Δ*copA*Δ*adcA,* and Δ*copA*Δ*cntA* mutants to Cu intoxication during growth in chelex-treated medium with or without CuSO_4_ supplementation. In the absence of CuSO_4_, all three strains grew similarly (Fig. 2A). Consistent with prior reports^28,31^, Δ*copA* was more susceptible to Cu than wild type (Fig. 2D-E). Notably, when supplemented with 1000 μM CuSO_4_, Δ*adcA* and Δ*copA* grew similarly to each other, with both having a defect relative to wild type (Fig. 2D-E). This suggests that, in isolation, reliance on Cnt-StP is as detrimental as loss of CopA when exposed to Cu stress. Furthermore, the use of the StP-Cnt system (Δ*copA*Δ*adcA*) enhanced the sensitivity of the Δ*cop*A mutant to Cu intoxication, whereas the use of the Adc system did not (Δ*copA*Δ*cntA*; Fig. 2B-E). The Δ*copA*Δ*adcA* mutant was profoundly sensitive, with the lowest concentration of CuSO_4_ tested, 15.6 μM, sufficient to suppress growth (Fig. 2B, E). Plasmid-based expression of AdcA or addition of 10 μM ZnSO_4_ abrogated the growth defects (Fig. 2F, Supplemental Fig. 2). These data indicate that the phenotypes were driven by Zn limitation and reliance on the Cnt-StP system, respectively. Following growth in a Zn-limited medium and plating onto agar containing 25 μM CuSO_4_, Δ*copA* was 10- to 100-fold more sensitive than wild type bacteria in 100% of assays (5/5). Forcing *S. aureus* to rely on the StP-Cnt system further sensitized the bacterium to Cu intoxication, as Δ*copA*Δ*adcA* was 10-fold more sensitive to Cu than Δ*copA* or Δ*copA*Δ*cntA* (Fig. 2G). Similarly, in USA300 LAC strains lacking both CopA and CopBL^28^, reliance on the Cnt system sensitized the bacterium to Cu 10-fold more than strains relying on the Adc system (Supplemental Fig. 3B).

As Cu negatively impacts both cellular processes and Zn uptake, we evaluated how the loss of Cu detoxification impacted these processes by assessing GAPDH activity and cellular metal accumulation (Fig. 3 & 4). In the absence of Cu, the CopA null derivatives of the wild type, Δ*adcA* and Δ*cntA,* had equivalent GAPDH activity (Fig. 4A). In the presence of 15 μM CuSO_4_, Δ*copA* and Δ*copA*Δ*adcA* showed significantly reduced GAPDH activity compared to wild type in the same medium or wild type and themselves with 0 μM CuSO_4_. Furthermore, Δ*copA*Δ*cntA* had greater enzyme activity by comparison to Δ*copA or* Δ*copA*Δ*adcA* in the presence of Cu. Collectively, these data indicate that StP-associated Cu uptake is a substantial source of cellular Cu that necessitates the use of the Cu stress response to maintain bacterial fitness. In the absence of Cu, the Δ*copA*, Δ*copA*Δ*adcA*, and Δ*copA*Δ*cntA* mutants accumulated Cu at levels similar to their respective *copA-*encoding parental strains (Fig. 3A). Following growth in the presence of 15 μM and 500 μM CuSO_4_, Δ*copA* and Δ*copA*Δ*adcA* accumulated ∼2-3-fold more ^63^Cu than the wild type and Δ*adcA*. By comparison to their growth in the unsupplemented medium, their relative ^63^Cu accumulation increased by ∼200-300-fold (Fig. 3C, E). Notably, in the StP-Cnt-dependent Δ*copA*Δ*adcA* mutant ^63^Cu accumulation was higher than in the Δ*copA* mutant. Similar to wild type and Δ*cntA*, following growth in 500 μM CuSO_4,_ there was no difference in ^63^Cu accumulation in Δ*copA* and Δ*copA*Δ*cntA* (Fig. 3E). The StP-Cnt-dependent Δ*copA*Δ*adcA* strain had reduced cellular ^66^Zn accumulation upon supplementation with 500 μM CuSO_4_, consistent with the Δ*adcA* strain (Fig. 3F), while increased ^55^Mn and ^60^Ni were observed (Supplemental Fig. 4).

### Nutritional immunity enhances the sensitivity of *S. aureus* to Cu intoxication

During infection, extracellular Zn limitation is imposed by the host protein calprotectin (CP), which can reach concentrations of more than 1 mg/ml at infection sites^32^. As CP-mediated Zn restriction can induce expression of the StP-Cnt system ^10^, we investigated if the metal withholding response of the host could sensitize *S. aureus* to Cu intoxication. Compared to a metal-replete medium, treatment with CP resulted in a lower concentration of Cu being necessary to induce *copA* expression (Fig. 5A). This indicates that CP-imposed Zn starvation could sensitize *S. aureus* to *in vivo* Cu intoxication.

**Figure 5.**
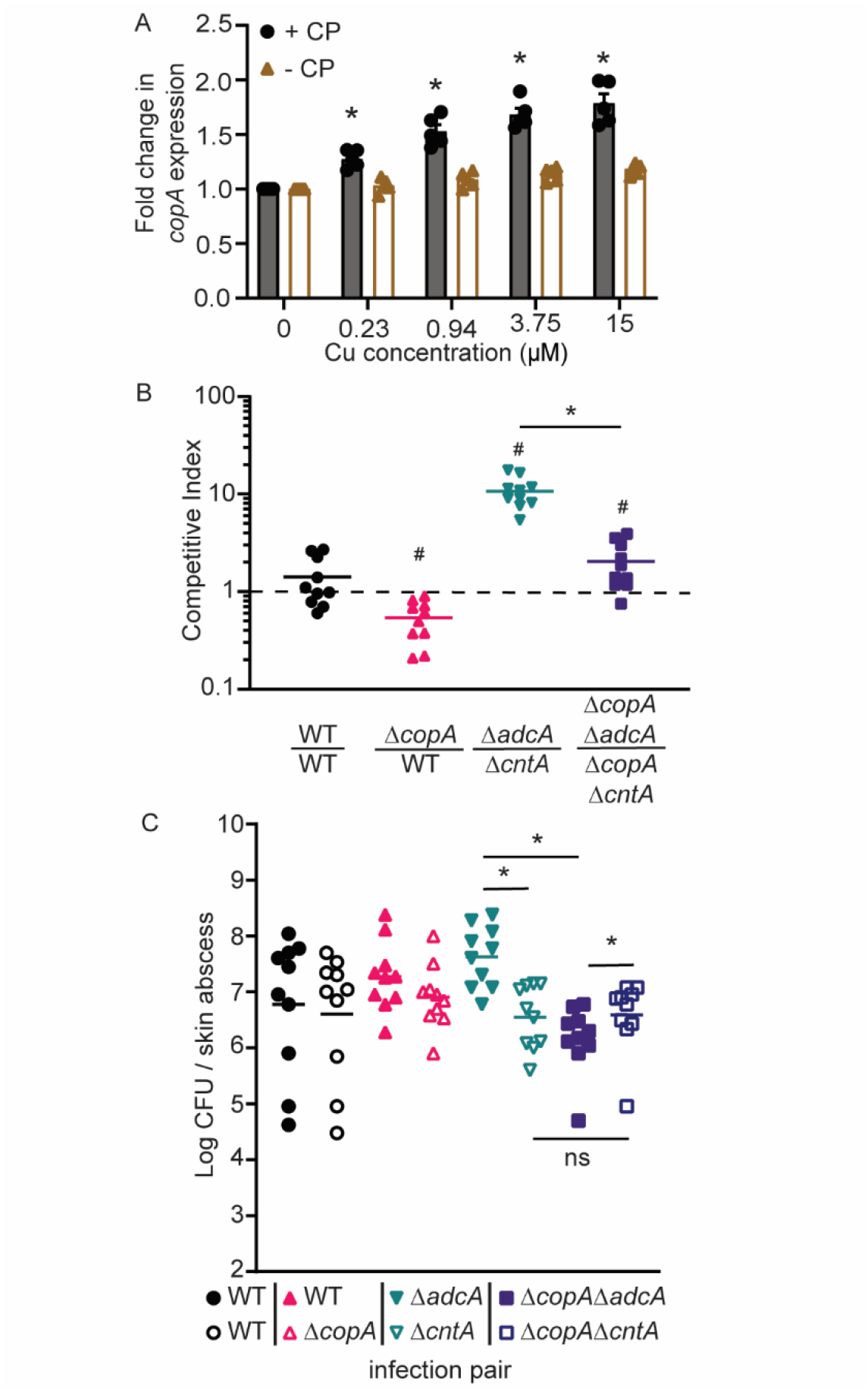
The Cnt system leads to copper import *in vivo*. (A) *S. aureus* Newman wild type carrying the P*_copA_*-YFP reporter was grown in metal-replete medium in the presence or absence of 960 μg/mL CP then exposed to a range of CuSO_4_ concentrations in Zn-limited medium. The expression of *copA* was assessed by measuring fluorescence at T = 4. * = p ≤ 0.05 relative to the same strain grown without CP via two-way ANOVA with Sidak’s multiple comparisons test. N = 5. Error bars = SEM. Wild type C57BL/6 were subcutaneously infected with equivalent CFUs of the indicated *S. aureus* pairs, and (B) competitive indices (CI), and (C) bacterial burdens were determined 7 days post-infection. (C) Closed and open symbols of the same color and shape denote the two mutants in each pair. The wild type vs. wild type comparison (circles) is a competition between wild type bacteria carrying the two different antibiotic cassettes used and shows that they do did not alter the outcome of infection. (B & C) * = p < 0.05 for the indicated comparisons via Mann-Whitney U test, # = p ≤ 0.05 compared to a theoretical mean of 1 via One sample t-test. n = 10.

### The use of staphylopine leads to copper import during infection

While phagocytes are known to impose Cu intoxication on *S. aureus* ^17,26^, the tissues in which *S. aureus* experiences Cu intoxication during infection remain unknown. This limits investigations into how Cu intoxication protects the host from *S. aureus* infections and the subversion mechanisms employed by the bacterium. *S. aureus* is a frequent cause of skin infections^33,34^. Accordingly, the potential for Cu intoxication to contribute to infection control during skin infection was investigated. Using a competition model of subcutaneous infection, the ability of Δ*copA* to cause infection was compared to wild-type bacteria. After 7 days of infection, loss of CopA diminished the ability of *S. aureus* to cause infection (Fig. 5B). After 14 days, a competitive index could not consistently be calculated, as the Δ*copA* mutant could no longer be detected in 3 out of 5 mice used for pilot studies, while all mice remained infected with the wild type strain. These observations reveal that within the dermis, *S. aureus* must overcome host-imposed Cu intoxication to successfully cause infection.

Next, we sought to determine if the StP-Cnt system sensitizes *S. aureus* to Cu intoxication during infection by competing Δ*adcA* against Δ*cntA* and Δ*copA*Δ*adcA* against Δ*copA*Δ*cntA*. Consistent with the necessity of the StP-Cnt system for infection^10,35^, the StP-Cnt-dependent Δ*adcA* mutant out-competes the Δ*cntA* mutant. However, this competitive advantage is reduced when the ability to manage cellular Cu stress is compromised, as shown by the competition between the Δ*copA*Δ*adcA* and Δ*copA*Δ*cntA* mutants (Fig. 5B). Furthermore, the data show that Δ*copA*Δ*adcA* had a reduced bacterial burden compared to Δ*adcA* in skin abscesses (Fig. 5C). These results indicate that, during infection, Cu intoxication poses a substantial risk to strains using the Cnt-StP system relative to those reliant upon the Adc system. Collectively, this work shows that the limited selectivity of StP for divalent cations opens the way to cellular Cu accumulation during infection.

## DISCUSSION

Transition metals are crucial nutrients, but they can also be profoundly toxic, with both properties exploited by the host’s innate response to infection^7,17,36,37^. The current investigation revealed that host-imposed metal starvation enhances the efficacy of Cu intoxication, with Zn starvation reducing the Cu concentration necessary to activate the Cu stress response of *S. aureus* by 1000-fold. Further work revealed that StP and the Cnt importer enable Cu to gain access to the bacterial cytoplasm, and that import via this transporter can overwhelm the Cu detoxification machinery. A recent study revealed that during infection, Cu abundance increases ∼150-fold in Zn-depleted regions of tissue^38^. This leads to a model (Fig. 6) where metallophores are used to respond to a primary threat, Zn starvation, creating a second threat, Cu intoxication. The current observations also challenge the conventional dogma that the lack of specificity often associated with StP-like molecules and other metallophores is broadly beneficial^6,39^.

**Figure 6.**
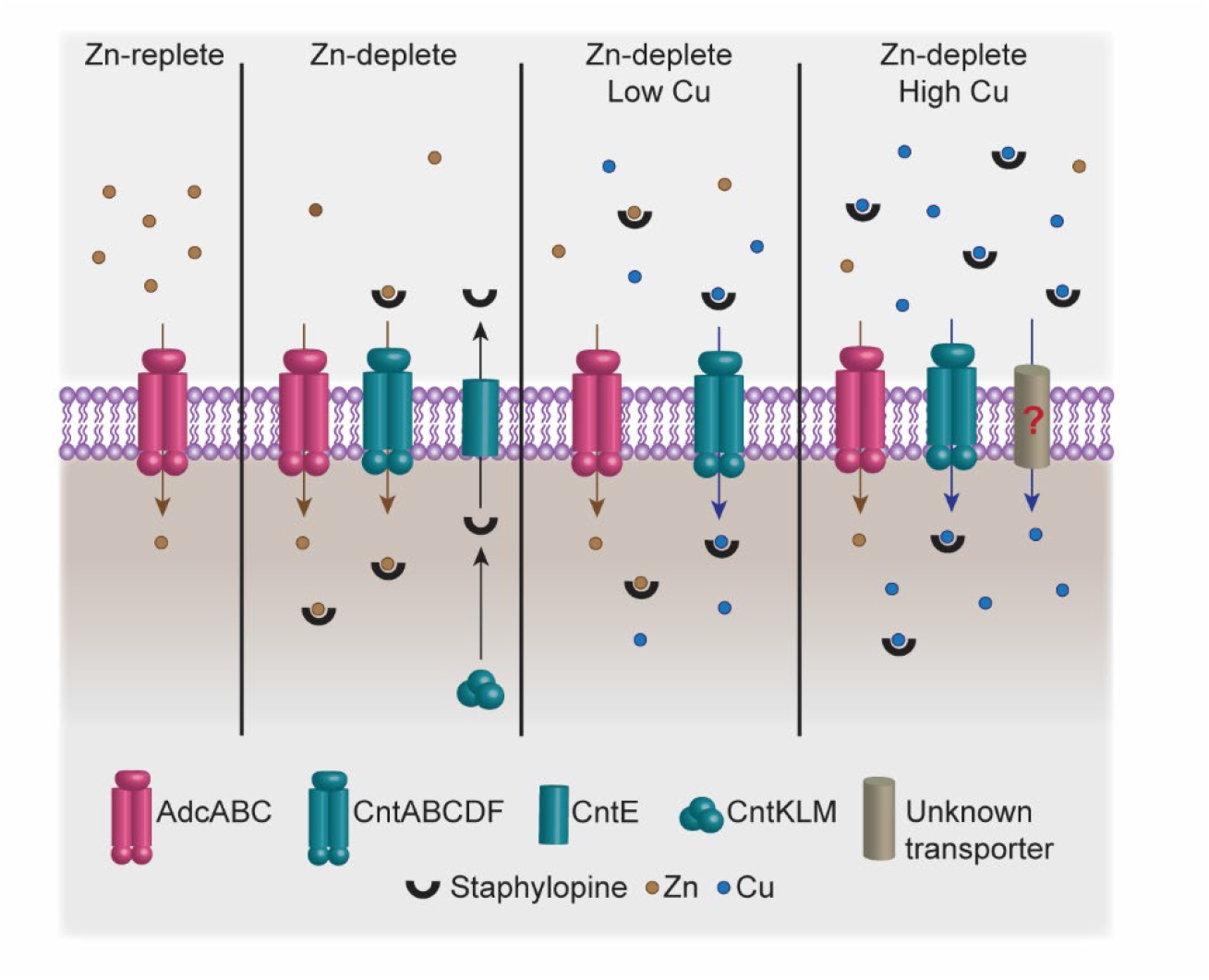
Model of Metallophore-driven Cu intoxication. In Zn-replete conditions, the classical AdcABC system is sufficient to meet cellular Zn requirements. In Zn-deplete environment, *S. aureus* employs the Cnt-StP system to acquire Zn. In the presence of low levels of Cu, StP binds and imports Cu, leading to cellular accumulation of Cu and induction of the Cu stress response. As concentrations increase, Cu blocks metallophore-dependent import of Zn and gains access to the cytoplasm via unknown additional mechanisms.

Cu detoxification systems are widespread in bacteria suggesting that Cu intoxication is a threat bacteria frequently encounter^21,40^. However, millimolar concentrations of Cu are frequently necessary to observe phenotypes under standard laboratory culture conditions^29,41^. This is substantially higher than the concentrations present in human tissues and fluids, which range from 0.1 to 20 μg/g in healthy individuals^42,43^. Even in Cu-rich environments, such as phagolysosomes, concentrations are generally in the micromolar range^42,44^. The observation that Zn limitation and calprotectin reduce the concentration of Cu necessary to intoxicate *S. aureus* resolves this disconnect. It also emphasizes the importance of considering the threats microbes face in their totality rather than in isolation.

*S. aureus* is not the only microbe in which StP-like molecules could drive copper accumulation. *Staphylococcus epidermidis* colonizes the skin, encodes an StP-Cnt system, and many isolates contain more than one Cu detoxification system^28,45^. In *P. aeruginosa,* PsP synthesis is enhanced in individuals with cystic fibrosis and is necessary to infect the lung, a tissue in which other pathogens must overcome Cu intoxication^42,46^. Similarly, CopA is necessary for *P. aeruginosa* infection of the liver, a tissue in which *S. aureus* relies on the StP-Cnt system to obtain Zn^22^. Environmental microbes also encounter Zn-limited niches^47,48^. Accordingly, genomic analysis suggests that actinobacteria, firmicutes, proteobacteria, and fusobacteria utilize opine-metallophores^2,3^. Cu detoxification systems are also widespread in environmental microbes, regularly co-occurring with StP-like metallophore synthesis and transport genes^49^. Environmental microbes must also contend with predation by amoeba, which use Cu to kill phagocytosed organisms^50^. Taken together, these observations suggest that opine-type metallophores may open the way to Cu intoxication in both infectious and environmental microbes. The threat of intoxication created by StP-like molecules may not be limited to Cu. In addition to Cu, opine metallophores can chelate multiple metals, including nickel, cobalt, cadmium, and lead, to which environmental microbes are potentially exposed^15,51^. Cumulatively, these observations support a model wherein StP-like metallophores may be an Achilles’ heel that renders invading pathogens susceptible to host-mediated metal intoxication.

Recently, it has become apparent that siderophores, which classically contribute to Fe(III) acquisition, can contribute to divalent cation acquisition, with yersiniabactin being necessary for *E. coli* and *Y. pestis* to compete with CP for Zn and obtain this metal within the host^52,53^. In Uropathogenic *E. coli* (UPEC), yersiniabactin also binds and imports Cu into the periplasm and cytoplasm. This activity is thought to benefit UPEC^6,39^. However, UPEC encounters Cu toxicity during urinary tract infections, and the Cus system, which removes Cu from the cytoplasm, is necessary for infection^54,55^. Furthermore, as with most bacteria, the mechanism by which toxic Cu concentrations gain access to the cytoplasm is unknown. The current investigations suggest that Zn limitation and subsequent use of yersiniabactin may also drive toxic Cu accumulation. Notably, yersiniabactin and the machinery necessary to import it in complex with divalent cations are present in other pathogenic *E. coli* as well as *Yersinia* and *Klebsiella* species^52,53,56^.

The significance and biological impact of non-cognate metal-binding by metallophores has not previously been defined. While StP, and by extension related metallophores, provide crucial benefits to pathogens at the host-pathogen interface, this work establishes that the promiscuity of these small molecules can also render microbes susceptible to metal intoxication.

## MATERIAL AND METHODS

### Ethics Statement

All experiments involving animals were approved by the Institutional Animal Care and Use Committee of the University of Illinois at Urbana-Champaign (IACUC license number 20257) and performed according to NIH guidelines, the Animal Welfare Act, and US federal law.

### Bacterial Strains and Growth Conditions

*S. aureus* Newman and LAC derivatives were used for all experiments (Table 1). *S. aureus* strains were grown at 37 °C in either 5 mL tryptic soy broth (TSB) on a roller drum or on tryptic soy agar (TSA) plates for performing routine culturing or genetic manipulation. *E. coli* strains were routinely cultivated at 37 °C in Luria broth (LB) with shaking or on Luria agar plates. For plasmid maintenance in *E. coli* and *S. aureus*, when appropriate, antibiotics were added at the following final concentrations: 100 μg/mL ampicillin, 50 μg/mL kanamycin, 10 µg/mL trimethoprim, 10 µg/mL chloramphenicol, and 1 μg/mL tetracycline. Both bacterial species were stored at −80 °C in a growth medium containing 30% glycerol. The Newman Δ*adcA,* Δ*cntA*, Δ*cntKLM*, *ΔcopA,* Δ*copA*Δ*adcA,* Δ*copA*Δ*cntA,* Δ*zur*, Δ*zur*Δ*adcA,* and Δ*zur*Δ*cntA* strains were generated as previously described^10,57^. Briefly, the 5’ and 3’ flanking regions of the genes were amplified using the indicated primers (Table 3). They were then cloned into pKOR1, and the deletion was generated via allelic replacement^58^. The LAC *adcA::Tn, cntA::Tn, ΔcopAZ ΔcopBL adcA::Tn,* and *ΔcopAZ ΔcopBL cntA::Tn* strains were generated as previously described^59^. For complementation constructs, the *adcA*, *cntA* and *copA* coding sequences were amplified using the indicated primers and cloned into either pOS1 under the control of the constitutive *lgt* promoter or pKK30 under the control of the native promoter or the *lgt* promoter (Supplementary Table 2)^60,61^. For the fluorescent reporters, the promoters of the *cnt* operon and *copA* were cloned into the yellow fluorescent protein (YFP)-containing vector pAH5^62^. All constructs were verified by sequencing, and all mutants were confirmed to be hemolytic. See Table 1 and 2 for a complete list of the strains and plasmids used in this study.

**Table 1:**
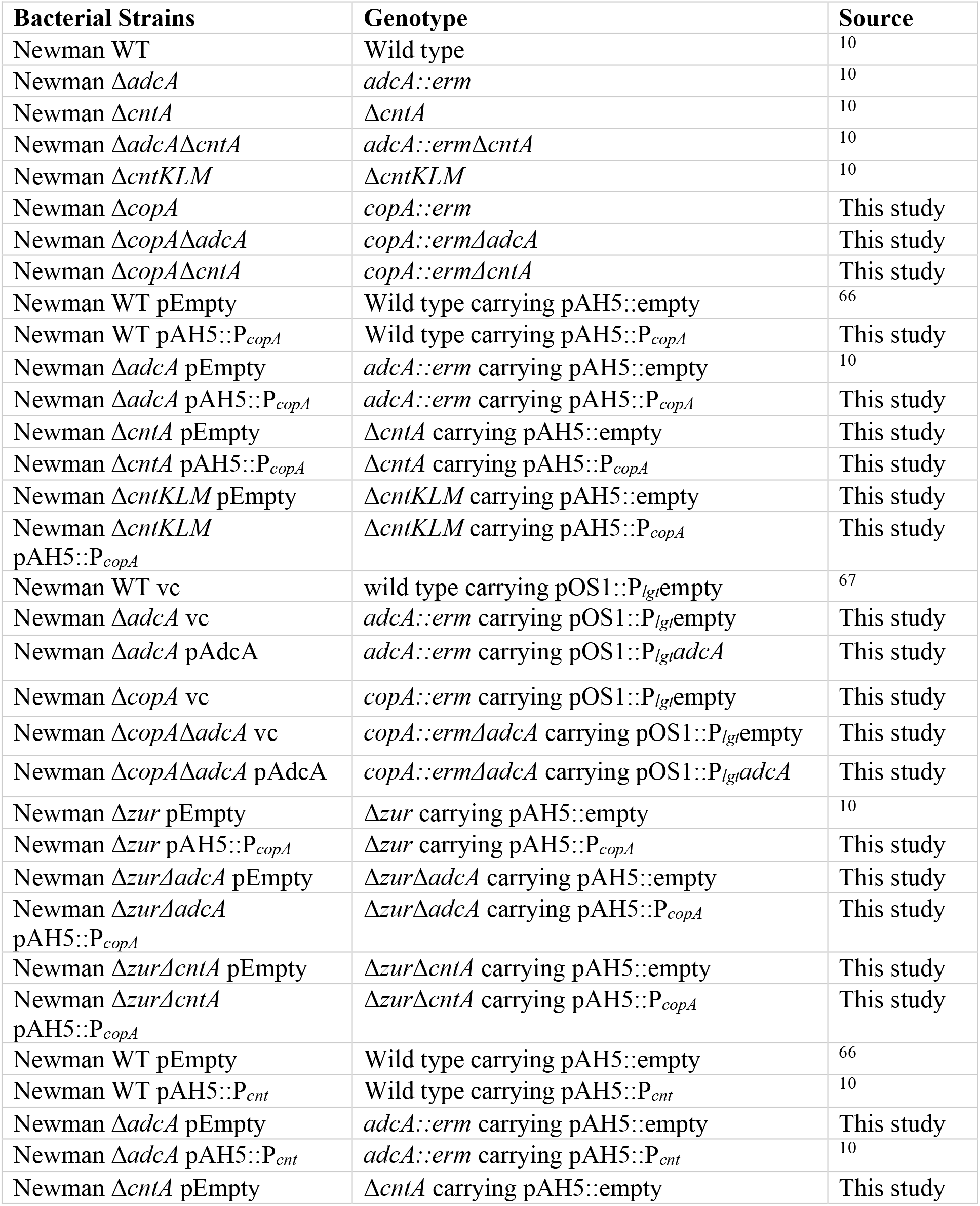
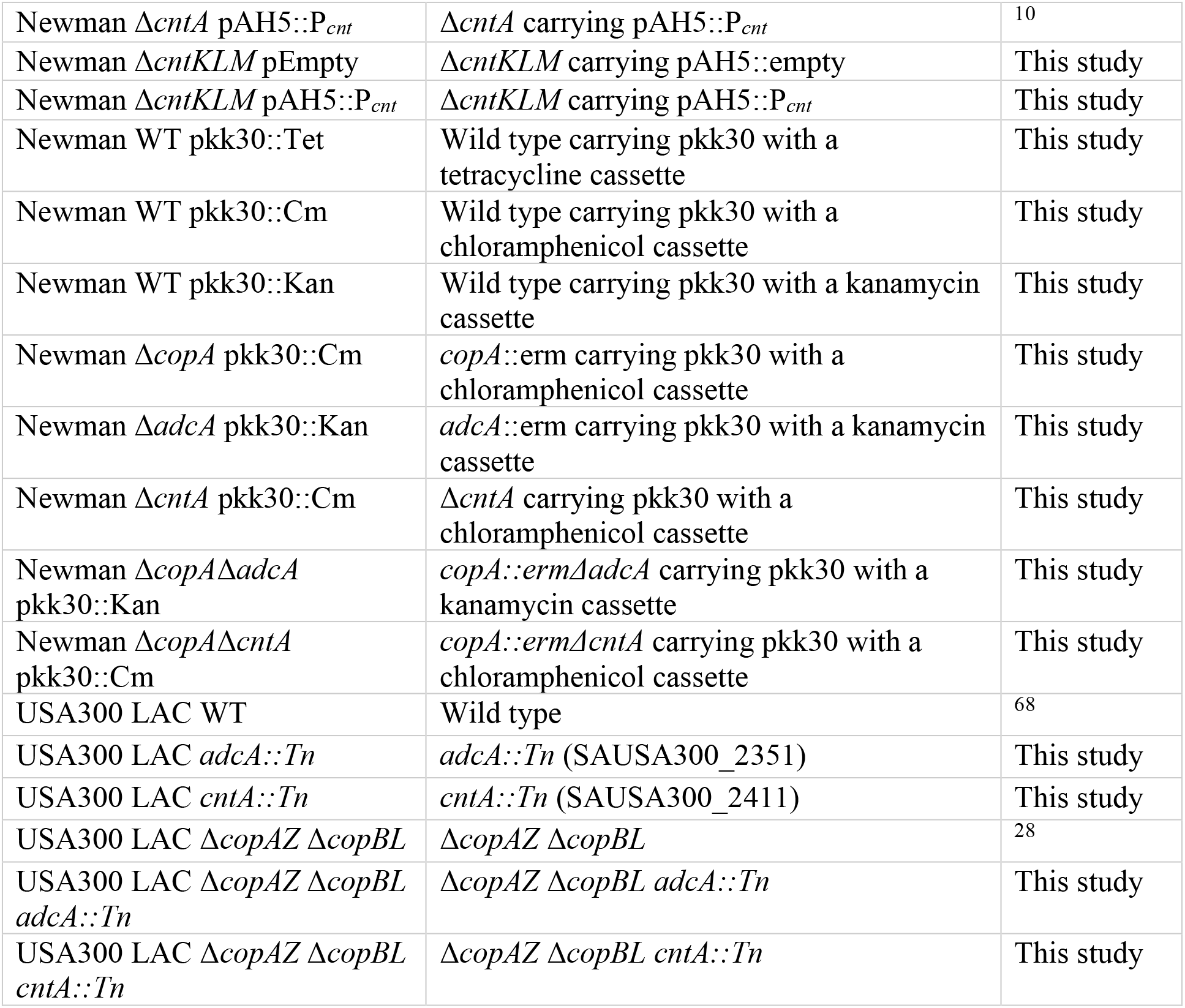
*Staphylococcus aureus* strains used in this study.

**Table 2:** Plasmids used in this study.

| Plasmid | Description | Reference |
| --- | --- | --- |
| pAH5::empty | pAH5 lacking a promoter for the expression of YFP | 62 |
| pAH5::P<sub>copA</sub> | Plasmid for *copA* promoter dependent YFP expression | This study |
| pAH5::P<sub>cnt</sub> | Plasmid for *cnt* operon promoter dependent YFP expression | 10 |
| pOS1::P<sub>lgt</sub>empty | complementation plasmid containing the *lgt* promoter | 69 |
| pOS1::P<sub>lgt</sub>*adcA* | complementation plasmid expressing AdcA under the control of *lgt* promoter | 10 |
| pkk30::Tet | Stable plasmid with a chloramphenicol cassette | This study |
| pkk30::Cm | Stable plasmid with a tetracycline cassette | This study |

**Table 3:**
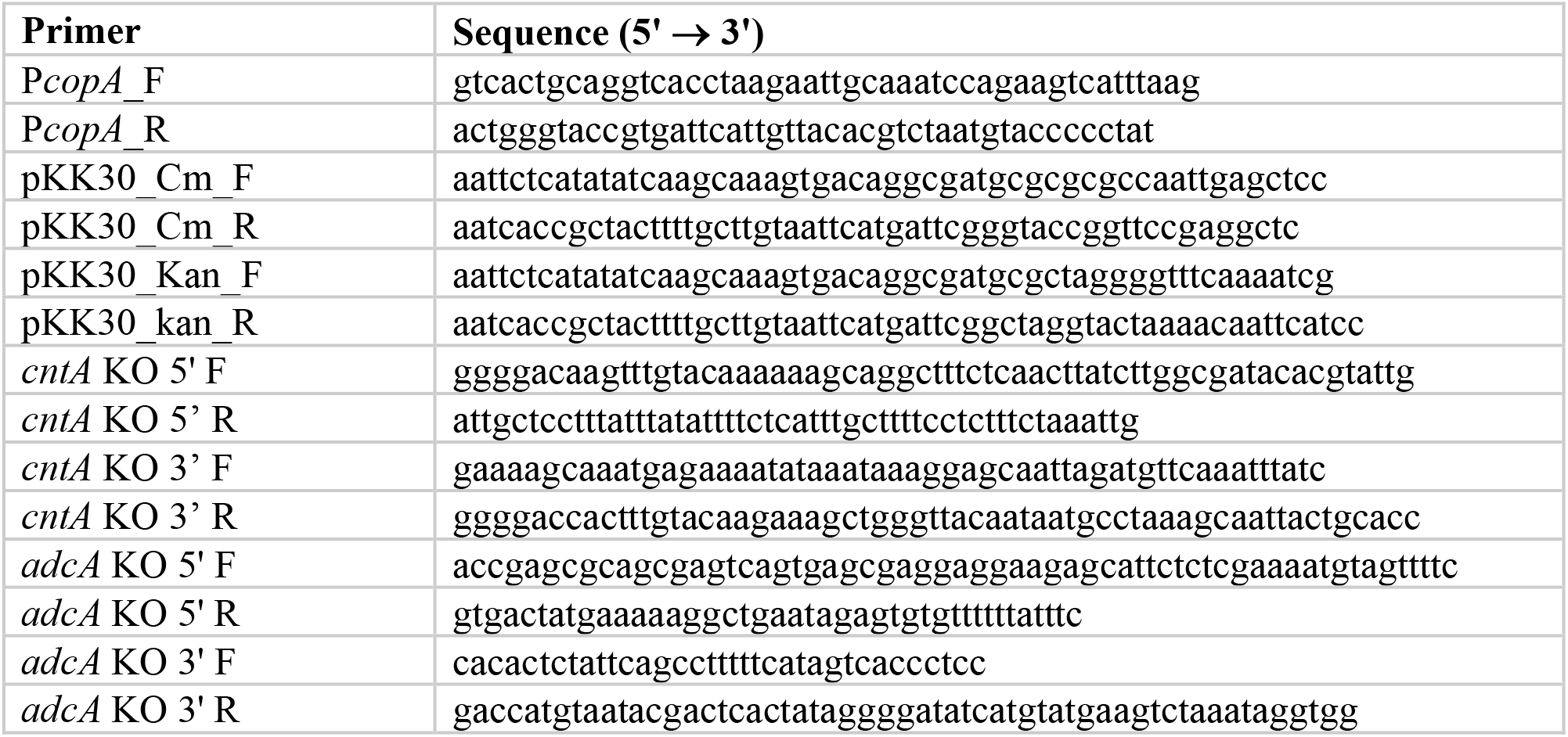
Primers used in this study.

### Expression Analyses

To assess the expression of *copA* and the *cnt* operon, *S. aureus* strains were grown overnight in chelex-treated RPMI plus 1% casamino acids (NRPMI) supplemented with 1 mM MgCl_2_, 100 µM CaCl_2_, and 1 µM MnCl_2 27_, and 10 μg/mL of chloramphenicol. Next morning, the overnight cultures were diluted 1:100 in 100 μL of the same growth medium with or without 10 μM ZnSO_4_ in a 96-well round-bottomed microtiter plate and grown to mid-log phase (about 4 hours) with orbital shaking (180 rpm) at 37 °C. For assays using CP to impose metal limitation, the overnight cultures were diluted 1:100 in 96-well round-bottom plates containing 100 μL of a growth medium, which consisted of 38% TSB and 62% calprotectin buffer (20 mM Tris, pH 7.5, 100 mM NaCl, and 3 mM CaCl_2_) in the presence or absence of 960 μg/mL of CP. For both assays using metal-defined medium and CP, the bacteria were harvested and resuspended in fresh NRPMI supplemented with 1 mM MgCl_2_, 100 µM CaCl_2_, and 1 µM MnCl_2_ at equivalent ODs. The bacteria were then diluted five-fold in 100 μL NRPMI supplemented with 1 mM MgCl_2_, 100 µM CaCl_2_, 1 µM MnCl_2_ and 10 μg/mL of chloramphenicol in the presence or absence of 10 μM of ZnSO_4_ and various CuSO_4_ concentrations in 96-well round-bottomed microtiter plates. The bacteria were incubated with orbital shaking (180 rpm) at 37 °C, and growth was measured by assessing optical density (OD_600_) and fluorescence (Excitation: 505 nm, Emission: 535 nm) was assessed every hour using a BioTek Synergy H1 microplate reader.

### Copper Susceptibility Assays

For liquid culture assays, overnight cultures were grown in 5 mL TSB in 15 mL conical tubes at 37 °C on a roller drum. The overnight cultures were pelleted and resuspended in 5 mL NRPMI supplemented with 1 mM MgCl_2_, 100 µM CaCl_2_, and 1 µM MnCl_2._ They were then diluted 1:100 in 100 μL NRPMI supplemented with 1 mM MgCl_2_, 100 µM CaCl_2_, 1 µM MnCl_2,_ and a range of CuSO_4_ concentrations and 100 µM ZnSO_4_ as indicated. Bacteria were incubated with orbital shaking (180 rpm) at 37 °C, and growth was measured by assessing optical density (OD_600_) every 1-2 h using a BioTek Synergy H1 microplate reader. For spot plating assays using defined medium, the strains were grown overnight in NRPMI supplemented with 1 mM MgCl_2_, 100 µM CaCl_2_, and 1 µM MnCl_2 27_. The next morning, the cultures were diluted 1:100 in 100 μL of the same growth medium with or without 10 μM ZnSO_4_ in a 96-well round-bottomed microtiter plate and cultured to an OD_600_ of ∼0.5 with orbital shaking (180 rpm) at 37 °C. Following growth to an OD_600_ of ∼0.5-0.6, cultures were serial diluted and spot plated on RPMI Medium 1640 (Gibco) agar (1%) plates containing various Cu concentrations.

### Elemental Analyses

The elemental content of the *S. aureus* Newman strains was determined by ICP-MS essentially as previously described^63,64^. Succinctly, the strains were grown using the culturing parameters described for the copper susceptibility assays. Bacteria were harvested during log-phase growth (OD_600_ of ∼0.2) by centrifugation at 2,500 × *g* for 10 min and washed two times with 0.1 M ethylenediaminetetraacetic acid (EDTA) and then washed two further times with MilliQ water to remove adventitious trace element content. The cells were then suspended in 1 mL of MilliQ water, and an aliquot was collected for CFU determination. The bacteria were then centrifuged, and the supernatant was removed. Bacterial pellets were desiccated at 96 °C overnight and then weighed to determine the dry cell mass of the pellet. The cellular material was digested in 250 µL of 65% (v/v) HNO_3_ at 96 °C for 20 min. Insoluble material was removed by centrifugation at 20,000 × *g* for 25 min and the supernatant was diluted in MilliQ-H_2_O to a final volume of 1 mL and analyzed in technical triplicate by ICP-MS on an Agilent 8900 ICP-MS/MS.

### Glyceraldehyde-3-Phosphate Dehydrogenase Activity Assay

For glyceraldehyde-3-phosphate dehydrogenase (GAPDH) assays, *S. aureus* Newman strains were grown using the culturing parameters described for the copper susceptibility assays and harvested during log phase (OD_600_ ∼0.2). The cells were washed once with 50 mM sodium phosphate buffer (pH 7.5) containing 5 mM EDTA and then washed twice with 50 mM sodium phosphate buffer (pH 7.5) without EDTA. The cells were then resuspended in 500 μL of 50 mM sodium phosphate buffer (pH 7.5) and homogenized twice in a FastPrep-24 Bead beater at 6 m/s for 45 sec with 5 min of incubation on ice in between. Cell lysates were centrifuged at 4 °C in a microcentrifuge at 13,000 × g for 15 min. The protein concentration in the cell lysate was determined via BCA assay (Pierce). GAPDH activity in cell lysates was determined by adding 100 μL of cell lysate (containing 1 μg of total protein) to 100 μL of assay buffer (50 mM sodium phosphate buffer pH 7.5, 10 mM EDTA, 80 mM triethanolamine, 4 mM glyceraldehyde-3-phosphate (G3P; Sigma-Aldrich), 4 mM nicotinamide adenine dinucleotide (NAD^+^; Sigma-Aldrich) in a flat-bottomed 96-well plate. The reaction was followed by measuring absorbance at 340nm every 2 min for 30 min on a BioTek Synergy H1 microplate reader. Negative controls (lacking either G3P, NAD^+^, or cell lysate) were included in the assay.

### Animal Infections

All animal experiments were performed as previously described^33,65^. *S. aureus* strains were grown overnight in 5 mL TSB in 15 mL conical tubes at 37 °C on a roller drum. The overnight cultures were diluted 1:50 in fresh 10 mL TSB in 50 mL conical tubes and grown to early log phase (OD_600_ ∼ 0.4) at 37 °C in a shaking incubator at 180 rpm. Bacteria were centrifuged at 4000 rpm for 10 min and resuspended in phosphate-buffered saline (PBS) to a concentration of 1 × 10^9^ CFU/mL. Each bacterial competing pair, containing distinct antibiotic markers, were mixed in a 1:1 ratio and placed on ice. The flanks of ten-week-old female C57BL/6 mice were shaved and depilated with Nair before infection. Mice were injected subcutaneously with 50 μL of 1 × 10^9^ CFU/mL of *S. aureus* mutant pairs (containing either kanamycin or chloramphenicol antibiotic markers) as specified. The infection was allowed to proceed for seven- or fourteen-days, after which the mice were sacrificed. The skin abscesses were harvested and homogenized in phosphate-buffered saline (PBS). Bacterial burdens were enumerated by plating serial dilutions on appropriate antibiotic-containing TSA plates. Competitive indices (CI) were calculated. CI is defined as the bacterial output ratio divided by the bacterial input ratio used to initiate the infection.

## Statistical Analyses

All statistical analyses were performed using GraphPad Prism version 9 and the indicated statistical tests.

## Acknowledgments

This work was supported by the grants from The Vallee Foundation and US National Institutes of Health (R01AI155611 and R01AI118880) to TKF, the Australian Research Council Discovery Project (DP220100713) to CAM, the National Health and Medical Research Council Ideas Grant (2010400) to CAM, and NIAID grant 1R01AI139100-01 and USDA MRF project NE-1028 to JMB. SH was supported by Chester W. and Nadine C. Houston Endowment Fellowship, and SLN was supported by a Passe and Williams Fellowship. The views expressed in this work do not necessarily reflect those of the funders.

## Supplemental Figures

**Supplemental Figure 1:**
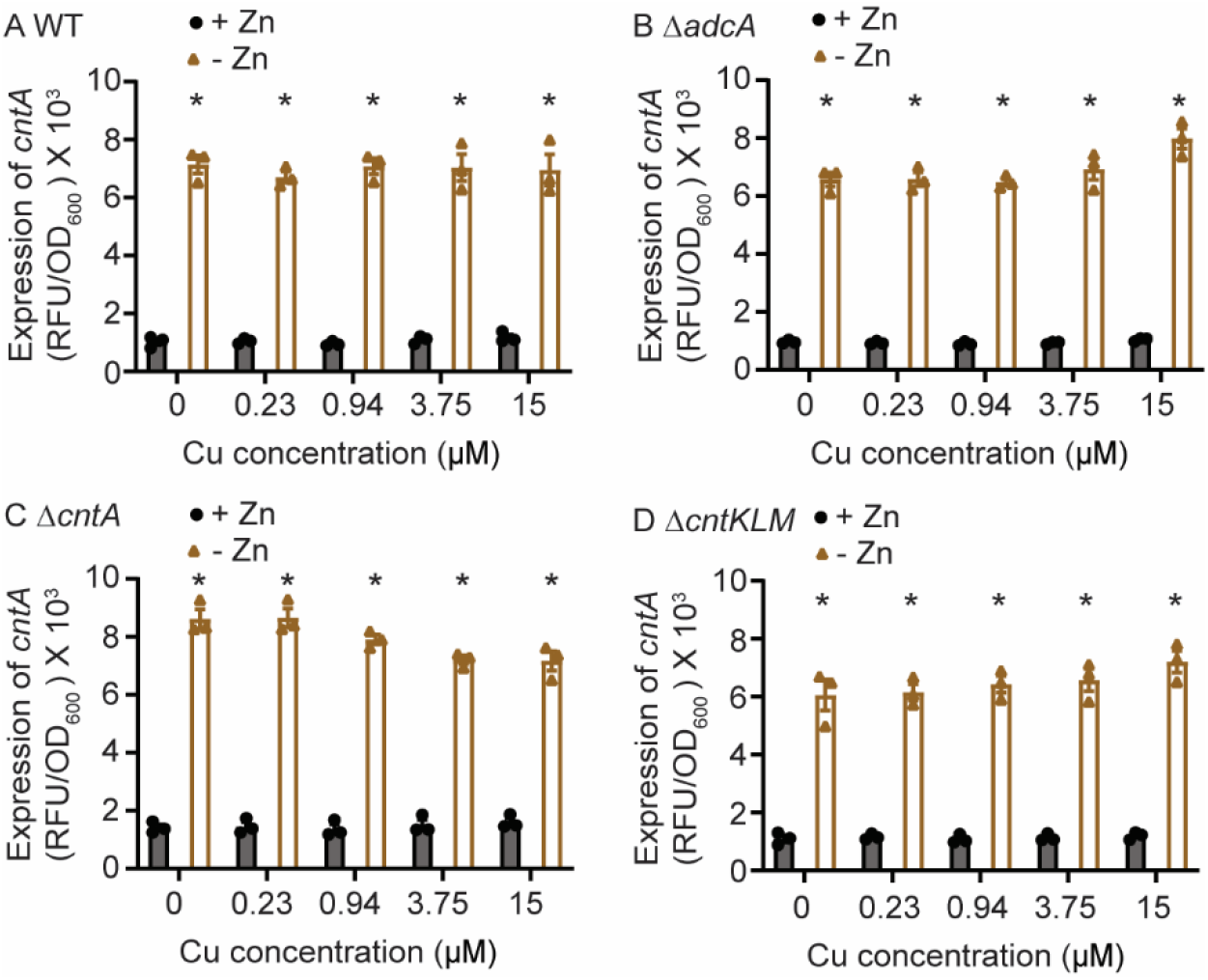
Zn limitation induces expression of the Cnt system in a Cu-containing medium. (A-D) *S. aureus* Newman wild type and the indicated strains containing P*_cnt_*-YFP reporter were grown in NRPMI containing a range of CuSO_4_ concentrations in the presence or absence of 10 μM ZnSO_4_ as specified. The expression of *cnt* was assessed by measuring fluorescence at T = 6. * = p ≤ 0.05 relative to the same strain with Zn via two-way ANOVA with Sidak’s posttest. n ≥ 3. Error bars = SEM.

**Supplemental Figure 2:**
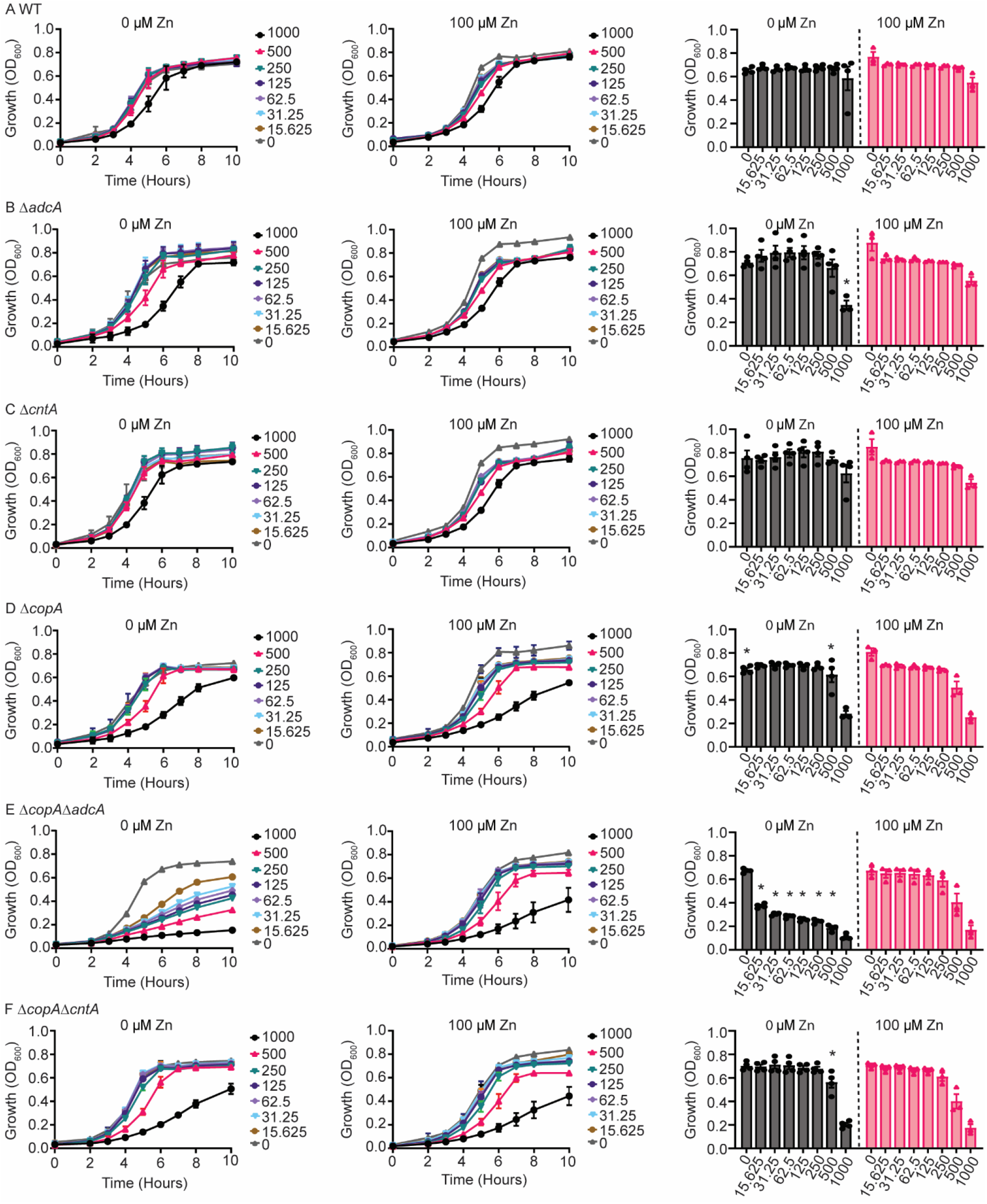
The Cnt system increases the susceptibility of *S. aureus* to Cu poisoning. (A-F) *S. aureus* Newman wild type and the indicated mutants were grown in the absence or presence of 100 μM ZnSO_4_ in NRPMI medium in the presence of various concentrations of CuSO_4_. Growth was assessed by measuring OD_600_ over time. Growth at 6 hours is shown and statistical analysis is present in the bar graphs. * = p ≤ 0.05 relative to the strain cultured under the same Cu concentration with Zn via two-way ANOVA with Sidak’s multiple comparisons test. Relevant statistical differences are being shown. n ≥ 3. Error bars indicate SEM.

**Supplemental Figure 3:**
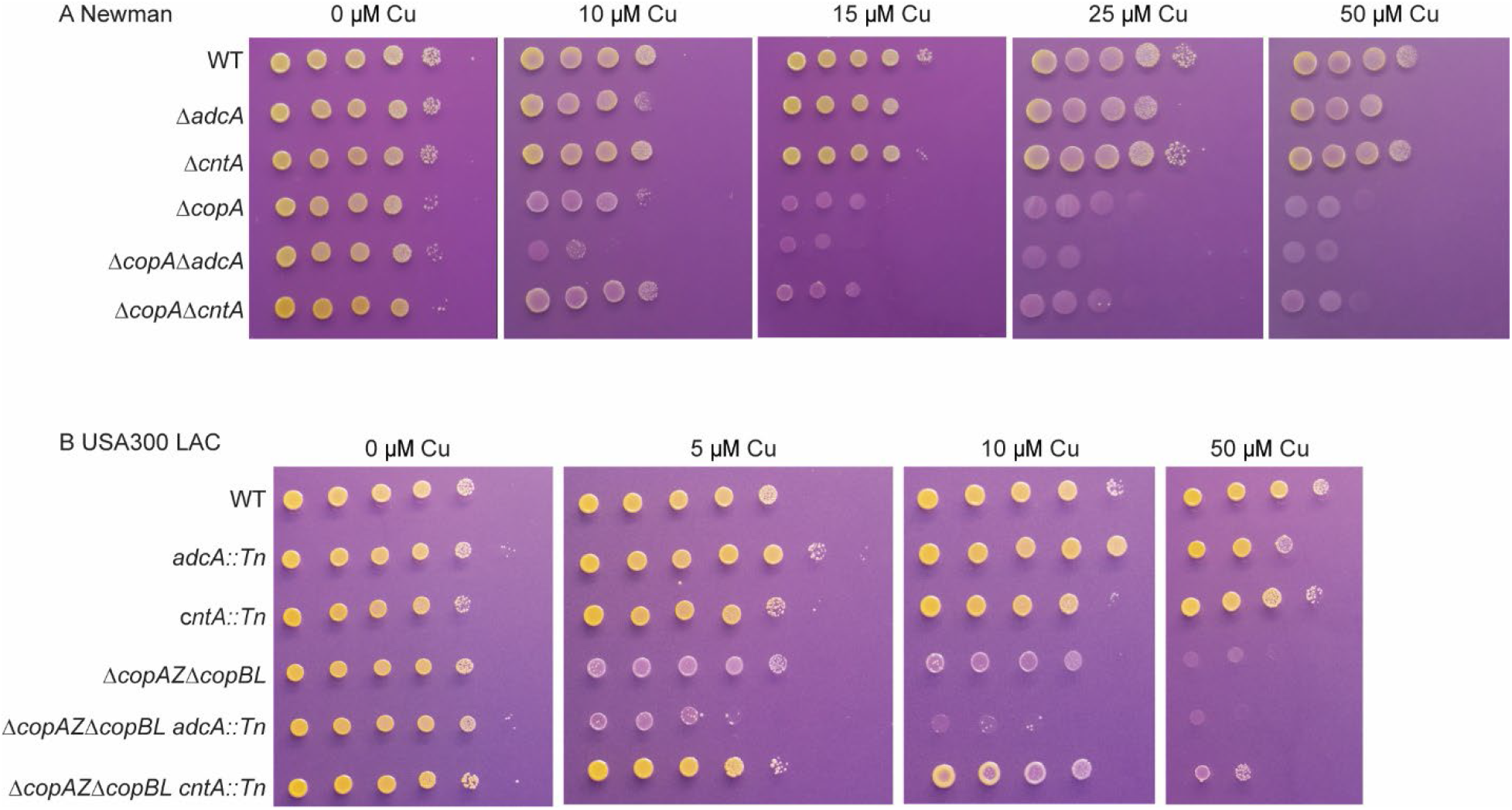
Reliance on the Cnt system increases Cu sensitivity in both Newman and USA300 LAC strain backgrounds. *S. aureus* (A) Newman and (B) USA300 LAC wild type strains and the indicated mutants were cultured in Zn-limited NRPMI and then spot plated onto plates with or without Cu as specified. Representative images of the spot plates are shown here.

**Supplemental Figure 4:**
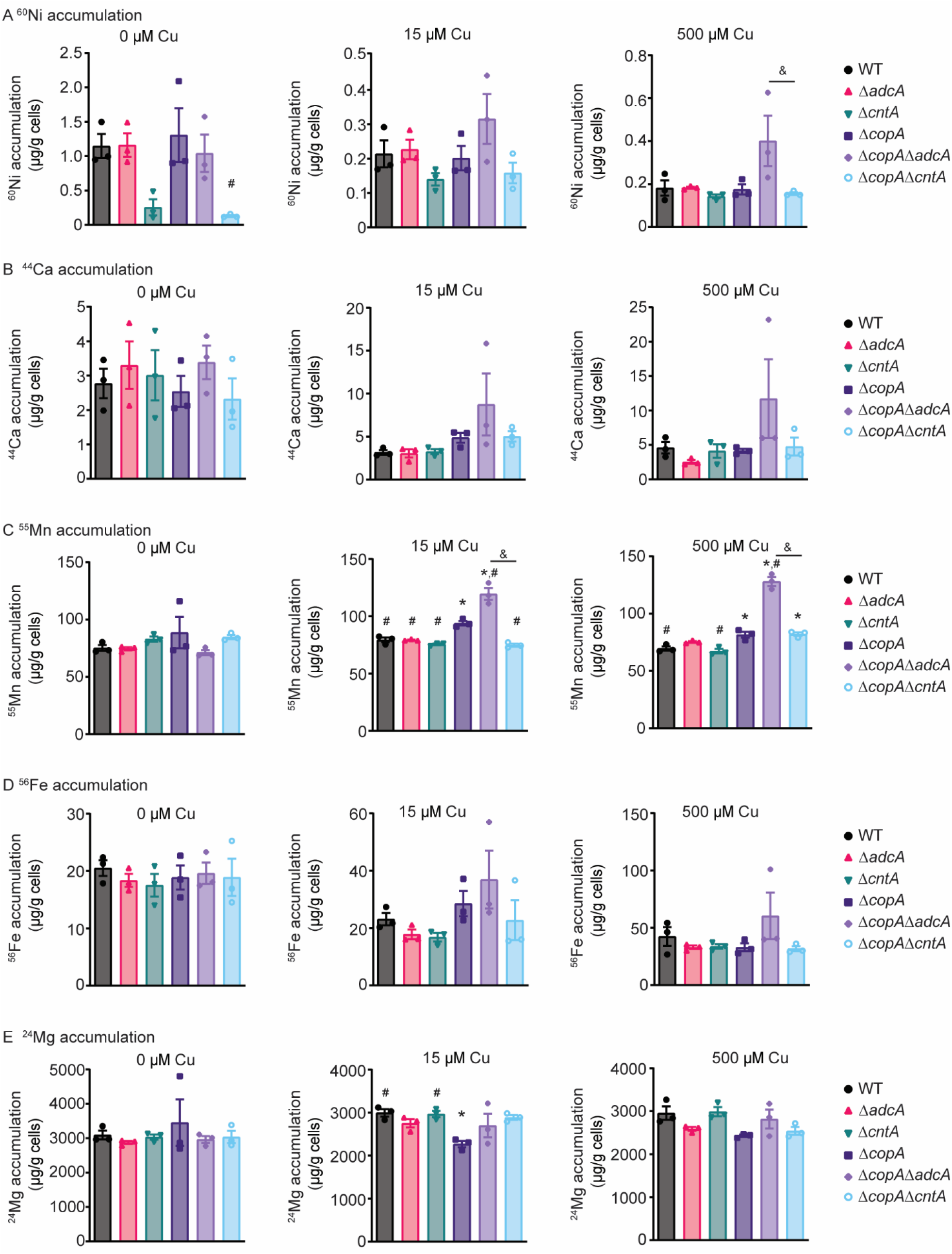
Metal accumulation in *S. aureus* when grown in the presence of Cu. *S. aureus* Newman wild type and the indicated mutants were grown in Zn-limited medium supplemented with 0 μM, 15 μM, and 500 μM CuSO_4_ and cellular (A) nickel (^60^Ni), (B) calcium (^44^Ca), (C) manganese (^55^Mn), (D) iron (^56^Fe) and (E) magnesium (^24^Mg) were assessed using ICP-MS. * = p < 0.05 via one-way ANOVA relative to wild type bacteria using Tukey’s posttest. # = p < 0.05 via one-way ANOVA relative to Δ*copA* using Tukey’s posttest. & = p < 0.05 via one-way ANOVA for the indicated comparison via Tukey’s posttest. n = 3. Error bars indicate SEM.

